# Limitations of fluorescent timer protein maturation kinetics to isolate transcriptionally synchronized cortically differentiating human pluripotent stem cells

**DOI:** 10.1101/2023.08.04.552012

**Authors:** Manuel Peter, Seth Shipman, Jeffrey D. Macklis

**Affiliations:** Department of Stem Cell and Regenerative Biology, and Center for Brain Science, Harvard University, Cambridge, MA, USA; Gladstone Institute of Data Science and Biotechnology, and Department of Bioengineering and Therapeutic Sciences, University of California, San Francisco, USA

**Author notes:** Correspondence should be addressed to J.D.M.

## Abstract

Differentiation of human pluripotent stem cells (hPSC) into distinct neuronal populations holds substantial potential for disease modeling *in vitro*, toward both elucidation of pathobiological mechanisms and screening of potential therapeutic agents. For successful differentiation of hPSCs into subtype-specific neurons using *in vitro* protocols, detailed understanding of the transcriptional networks and their dynamic programs regulating endogenous cell fate decisions is critical. One major roadblock is the heterochronic nature of neurodevelopment, during which distinct cells and cell types in the brain and during *in vitro* differentiation mature and acquire their fates in an unsynchronized manner, hindering pooled transcriptional comparisons. One potential approach is to “translate” chronologic time into linear developmental and maturational time. Attempts to partially achieve this using simple binary promotor-driven fluorescent proteins (FPs) to pool similar cells have not been able to achieve this goal, due to asynchrony of promotor onset in individual cells. Toward solving this, we generated and tested a range of knock-in hPSC lines that express five distinct dual FP timer systems or single time-resolved fluorescent timer (FT) molecules, either in 293T cells or in human hPSCs driving expression from the endogenous paired box 6 (PAX6) promoter of cerebral cortex progenitors. While each of these dual FP or FT systems faithfully reported chronologic time when expressed from a strong inducible promoter in 293T cells, none of the tested FP/FT constructs followed the same fluorescence kinetics in developing human neural progenitor cells, and were unsuccessful in identification and isolation of distinct, developmentally synchronized cortical progenitor populations based on ratiometric fluorescence. This work highlights unique and often surprising expression kinetics and regulation in specific cell types differentiating from hPSCs.

## Introduction

Differentiation of human pluripotent stem cells (hPSC) into distinct neuronal populations holds substantial potential for disease modeling *in vitro*^1-5^. This could enable both elucidation of pathobiological mechanisms and screening of potential therapeutic agents. However, detailed understanding of the transcriptional networks and their dynamic programs regulating endogenous cell fate decisions is critical for successful differentiation of hPSCs into subtype-specific neurons. Understanding of human-specific versions and subtleties of these networks and dynamics with hPSCs is very limited compared with major fundamental understanding of parallel neuron subtype differentiation in mice that has been deeply investigated over the past two decades^6-9^.

One major roadblock to this understanding and thus molecular manipulation for more refined and precise subtype-specific differentiation of hPSCs is the heterochronic nature of neurodevelopment, during which distinct cells and cell types in the brain and during *in vitro* differentiation mature and acquire their fates in an unsynchronized manner, hindering pooled transcriptional comparisons^7,10,11^. Thus, to optimally understand and potentially control the sequential steps of cell fate acquisition and neuronal maturation, it is important to identify and analyze cells that share the same developmental stage. One potential approach would be to re-organize and conceptually translate chronologic time into linear developmental and maturational time.

Single cell RNA sequencing (scRNAseq) is used extensively to investigate cellular and developmental heterogeneity during neuronal differentiation *in vitro*^12^ and *in vivo*^13^. However, scRNAseq typically only captures a relatively small part of the transcriptome due to limited sequencing depth, thus typically only allowing classification based on relatively highly expressed transcripts, and likely not capturing diverse and progressive cell states in a population of cells. One potential solution is to purify many cells at the same developmental– thus transcriptional– stage, enabling much deeper sequencing at sequential development stages to elucidate dynamics and regulation of gene expression.

One promising approach toward this goal is the widely used approach of genetically encoded fluorescent protein (FP) reporters expressed from cell-fate determining and cell type-specific promoters, typically used for real-time observation of cell states^14-17^. However, such FPs are binary, so individual cells within a pool of cells that all express a given FP might have begun expressing that FP at substantially different times, thus not allowing synchronization and analysis of more transcriptionally homogeneous cells based on promoter onset. It is important to know not just whether a cell expresses a cell-type marker, but also to cluster cells based on the timing and initiation of a distinguishing promoter activity. This would offer potential to optimally analyze dynamics and regulation of transcriptional state.

To circumvent these challenges, specific FP variants with changing fluorescence spectra and intensities over relatively short and reproducible time periods – fluorescent timers (FTs) – can be used as a form of molecular clock^18-23^. These FTs display time-dependent spectral conversion from one color to another during maturation, e.g. green to red or blue to red. In theory, this might allow determination of timing of promoter activation based on ratiometric fluorescence of the two colors. However, due to their typically low fluorescence intensities, FTs have not been used widely in mammalian systems.

Alternatively, two spectrally distinct FPs with distinct maturation constants– e.g., a fast-maturing green FP and a slow-maturing red FP– might be used as ratiometric molecular clocks without such limits of intensity^1,24^. Both FPs might be expressed from the same promoter, and the distinct and reproducible maturation times of the green and red FPs might theoretically enable reliable ratiometric measurement and determination of timing of promoter onset. FTs expressed from one or more endogenous, fate-determining promoters, in combination with fluorescence-activated cell sorting (FACS)^25,26^, might theoretically enable purification of cells synchronized based on their ratiometric fluorescence. Recently, an approach of this type has been used to study enteroendocrine progenitor cells in the intestinal epithelium of mice^27^. It is not known whether FPs and/or FTs also might be used to investigate transcriptional dynamics of fate decision and maturation of hPSCs toward human neural progenitors and neurons in *vitro*. This could “translate” chronological time to developmental-maturational time, thus might enable reliable analysis of substantially more transcriptionally homogeneous cell states.

In the work presented here, we generated and tested a range of knock-in hPSC lines that express five distinct FPs/FTs from the endogenous paired box 6 (PAX6) promoter^28^, highly expressed by dorsal cerebral cortex (cortical) progenitors of all cortical projection neurons. We differentiated each of these engineered hPSC lines toward cortical progenitor identity, and validated that each hPSC line differentiated into populations of progenitors with high levels of PAX6 and FT expression. We report here the negative results that none of the tested FP/FT constructs were successful in identification of distinct progenitor populations based on ratiometric fluorescence, thus unsuccessful toward isolating distinct, developmentally synchronized cortical progenitors.

## Results

### SlowFT expression in 293T cells and neural progenitor cells

In a first set of experiments, designed to assess dynamics of one FT within cultured mammalian cells, we generated stable knock-in 293T cells expressing the monomeric, blue to red maturing, FT SlowFT^19^ from the doxycycline (Dox) inducible Tet-ON promoter.

We induced SlowFT expression with a 24h pulse of Dox, and measured fluorescence changes at specific times after induction using flow cytometry. We first detected a strong increase of blue fluorescence 6h post-induction. After 24h, we detected double positive blue and red cells exhibiting increased fluorescence intensity over time, with red fluorescence highest in cells that also exhibited the highest blue fluorescence, indicating proper maturation of the FT (Supplemental Figure 1 A). These experiments demonstrate that SlowFT is expressed and properly matures in human cells, thus can be used to track promoter onset based on ratiometric blue to red fluorescence measurements.

The transcription factor PAX6 is highly expressed by the early human neuroectoderm *in vivo* and *in vitro*, and represses pluripotency genes during differentiation from hPSC into dorsal neural progenitor cells, making the endogenous PAX6 promoter an especially useful candidate as the first fate-determining promoter to target with an FT^28^. We used the CrispR/Cas9^29^ system to integrate SlowFT downstream of the endogenous PAX6 gene using a P2A splice acceptor site^30^ (Supplemental Figure 2 A). This leaves the endogenous PAX6 gene intact, and results in SlowFT expression levels comparable to endogenous PAX6 RNA, thus appropriately reporting PAX6 promoter onset (Supplemental Figure 2 B). We differentiated PAX6-SlowFT iPSC clones into PAX6-expressing dorsal neural progenitor cells^31^, and analyzed their fluorescence daily from the start of neuronal differentiation at day 0 up to day 8, using flow cytometry. Uninduced PAX6-SlowFT iPSC did not exhibit detectable blue or red fluorescence. We started to detect a strong increase of blue fluorescence from day 2 onward. However, contrary to our results with 293T cells, we did not detect an increase of red fluorescence over the same period to day 8 (Supplemental Figure 1 B).

Because PAX6 expression is virtually absent on day 2 in this *in vitro* neuronal differentiation system, we wondered if the strong increase of blue fluorescence might be simply evolving autofluorescence during neuronal differentiation, so we analyzed WT iPSC and WT progenitor cells at day 8 (both lacking any FT construct). Indeed, we found that WT cells showed a strong blue shift, indicating that the increase of blue fluorescence from day 2 onward in PAX6-SlowFT engineered cells is autofluorescence, and not fluorescence of SlowFT protein (Supplemental Figure 1 C). This was further confirmed by confocal imaging of PAX6-SlowFT neural progenitor cells on day 12, revealing strong PAX6 protein expression, but no detectable blue or red fluorescence (Supplemental Figure 1 D). Finally, we tested whether SlowFT mRNA is present in PAX6-SlowFT progenitor cells by RT-PCR on WT and PAX-SlowFT iPSC and progenitor cells at day 8. We found that PAX6 was highly expressed in WT and PAX6-SlowFT progenitors, and we detected SlowFT only in PAX6-SlowFT progenitors, indicating that SlowFT RNA is expressed by PAX6-SlowFT neural progenitors, and increases together with PAX6 mRNA (Supplemental Figure 1 E). While we detected strongly increased SlowFT mRNA, we did not detect SlowFT fluorescence, indicating either sub-detection limit intensity or potentially incorrect folding and maturation of SlowFT in human neural progenitors. Additionally, the strong increase of blue autofluorescence detected during WT neural differentiation could further mask low-intensity SlowFT fluorescence. Together, these results demonstrate that SlowFT is not suitable for tracking PAX6 promoter activation in neural progenitor cells.

### TandemFP expression in 293T and neural progenitor cells

Monomeric FTs like SlowFT have the advantage that they are small and therefore easy to incorporate into expression systems, but they can suffer from low intensity when expressed from endogenous promoters. Alternatively, a combination of spectrally distinct, fast maturing and slow maturing, FPs can be used as a fluorescent timer pair (dual fluorescent timer). In this approach, the useful range of the dual FT is determined by the difference in maturation times of the two FPs^1^. When expressed from an endogenous promoter (e.g., PAX6, Fezf2^32^), ratiometric fluorescence measurements between the fast and slow maturing FP theoretically might be used to time promoter activation. This approach would enable the use of bright fluorescence molecules, and any spectral combination is theoretically possible if their maturation times are significantly different, thus potentially making this system more flexible compared to a monomeric FT, and more suitable in systems in which more than one FT is expressed sequentially from distinct promoters.

To test the feasibility of such a dual fluorescent timer approach to purify progenitors with synchronized developmental/transcriptional time based on PAX6 promoter activation, we generated a new fluorescent timer (TandemFP) consisting of a fusion of fast-maturing green fluorescent protein sfGFP^33^ and slow-maturing red fluorescent protein DsRed2^34^ (Supplemental Figure 2 B). We chose this pair because DsRed2 matures 28-times slower than sfGFP, theoretically offering a long enough time window to measure promoter activation based on ratiometric fluorescence.

We generated 293T cells with Dox-inducible TandemFP, and pulse-induced them to test whether TandemFP faithfully reports promoter activation. Six hours after Dox induction, we detected strongly increased green fluorescence with further increases over time. At 24h, two distinct populations emerged– one GFP-only and one GFP/DsRed2double positive. At 24h, almost all cells are GFP positive and do not further increase their green fluorescence intensity. At 72h, only GFP/DsRed double positive cells were detected. (Figure 1 A, Supplemental Figure 3 A). These results demonstrate that the fluorescence changes observed follow reported maturation times for these FPs, and that ratiometric green-to-red measurements can be used to time promoter activation– something not possible using a single fluorophore alone.

**Figure 1:**
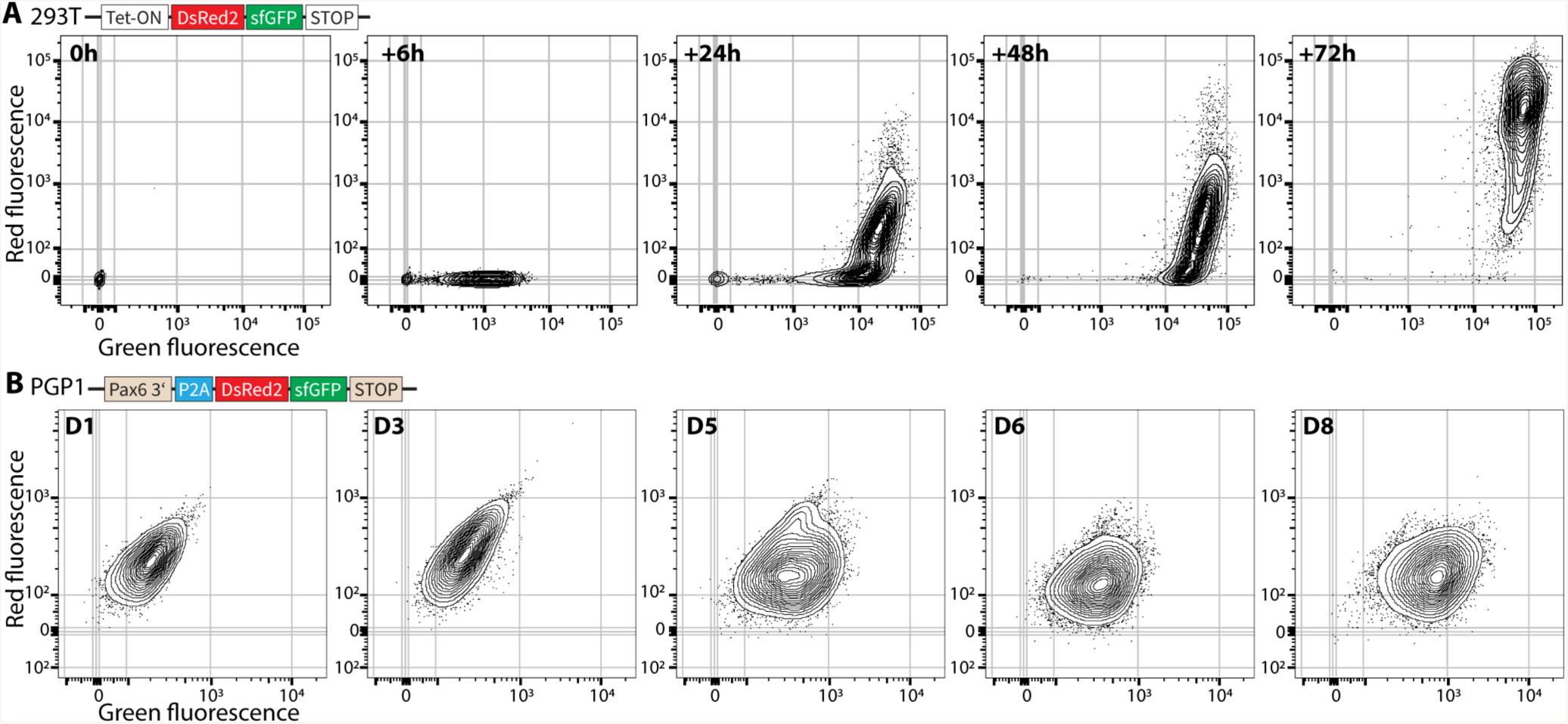
Distinct dynamics of TandemFP expression by 293T cells and human PGP-1 iPSC-derived neural progenitor cells. A: FACS plots of 293T cell expressing TandemFP from a Tet-ON promoter. Expression was induced by adding Dox to the cell culture medium. Cells were harvested and analyzed after 4 periods of culture– 6, 24, 48, and 72 hours (h). Green fluorescence increases strongly by 6h after induction, with most cells displaying both green and red fluorescence after 72 h. B: FACS plots of neural progenitor cells derived from PGP-1 iPSCs expressing TandemFP from the endogenous PAX6 promoter. PGP1-PAX6-tandemFP iPSCs were induced at day (D) 0, and neural progenitors were harvested and analyzed daily. Green fluorescence is detected by D5, however no red fluorescence is detected by D8.

To investigate whether TandemFP successfully times PAX6 promoter onset in human neural progenitor cells, we integrated TandemFP into the 3’ end of the PAX6 gene (Supplemental Figure 2 A, B). We differentiated these PAX6TandemFP iPSCs into neural progenitor cells, and longitudinally measured their fluorescence using flow cytometry. Based on our successful TandemFP experiments using 293T cells, we expected to first detect “GFP-only” progenitors, followed by GFP/DsRed2 double-positive progenitors, then a further increase of the DsRed2 signal.

We did not detect any fluorescence in PAX6-TandemFP iPSCs or neural progenitor cells until day 3 of differentiation. GFP-positive progenitors appeared from day 4 onward, with GFP fluorescence levels increasing until day 8. However, we did not detect GFP/DsRed2 double-positive progenitors (Figure 1 B). Further analysis on day 23 showed GFP single-positive but no double-positive cells.

These unanticipated results raised questions of whether the TandemFP fusion protein is correctly expressed and folded, or whether it does not properly mature in human neural progenitor cells. We investigated these questions by performing qPCR and western blot (WB) analysis in WT and PAX6-TandemFP expressing progenitor cells. We designed three qPCR primer sets that either target the sfGFP region, the DsRed2 region, or the middle region, covering both the sfGFP and the DsRed2 fusion region of the TandemFP mRNA. 12 days after induction, we detected strong upregulation, with all three primer sets in TandemFP progenitors, demonstrating that the TandemFP mRNA is fully expressed in neural progenitors. We further detected strong upregulation of PAX6 mRNA in both WT and TandemFP neural progenitors, indicating that the knock-in of TandemFP into the PAX6 locus did not alter its endogenous expression pattern (Supplemental Figure 3 A). Next, we investigated whether TandemFP mRNA is translated into a full-length fusion protein by performing WB against the N-terminal part of TandemFP using an antibody against GFP. Again, we found a strong upregulation of the PAX6 protein in both WT and TandemFP progenitors at day 8 of differentiation. When probing the same samples with the GFP antibody, we detected a strong band correlating with the molecular weight of TandemFP in the PAX6-TandemFP neural progenitors, indicating correct translation of the TandemFP DsRed2::sfGFP fusion protein. Interestingly we also detected a weaker band correlating with the MW of GFP (Supplemental Figure 3 C).

These results demonstrate that full length TandemFP is expressed and translated in human neural progenitor cells, and that its expression correlates with PAX6 mRNA and protein levels. However, we did not detect endogenous DsRed2 signal from the TandemFP fusion protein, indicating either issues with correct folding and maturation of the fusion protein or low DsRed fluorescence signal levels in progenitors, resulting in TandemFP not being suitable as a FT in human neural progenitor cells.

### Design and refinement of MolTimer dual fluorescent protein timer constructs

To address these suboptimal results in our next dual FP design, we aimed to increase brightness by using different FPs and adding a NLS^35^ to further increase signal by concentrating the FPs to the nucleus. We also separated the individual FPs with a P2A/T2A^30^ site, resulting in a single polycistronic mRNA that is translated into two separate FPs that are expected to fold and mature independently of each other. Recently, a very similar FT (Chrono)^27^ was successfully used to investigate enteroendocrine progenitor cells in the intestinal epithelium in mice. The Chrono design added a degron to the fast-maturing green fluorescent protein, thus increasing the effective time range of the FT. We based our next FT on a similar design, with some modifications. First, we changed dTomato^36^ to mScarlet^37^, which is roughly twice as bright as dTomato, with a slower maturation time, theoretically making it superior to dTomato in a FT construct. Second, we used a codonoptimized version of mNeonGreen (hmNeonGreen)^38^ to facilitate translation in human cells (Supplemental Figure 2 B).

We used this FT construct (MolTimer1.0) to generate knock-in iPSCs that express MolTimer1.0 from the endogenous PAX6 promoter. We differentiated these PAX6MolTimer1.0 iPSCs into neural progenitor cells, and longitudinally assayed MolTimer1.0 fluorescence in PFA-fixed cells using flow cytometry. Uninduced PAX6-MolTimer1.0 iPSCs did not exhibit fluorescence. From day 4 on, we detected a small population of red, mScarlet-positive cells that increased in both number and fluorescence intensity until day 10. They also exhibited weak, hmNeonGreen fluorescence. However, contrary to published maturation times that predicted faster maturation for mNeonGreen compared to mScarlet, we did not detect hmNeonGreenonly cells (Figure 2 A). These results were confirmed by longitudinal confocal imaging of endogenous mScarlet and hmNeonGreen fluorescence in PFA fixed PAX6MolTimer1.0 iPSCs and neural progenitor cells, in which we detected a linear increase of fluorescence for both FPs, but no hmNeonGreen-only cells (Figure 2 B). These results were not due to an incomplete expression of MolTimer1.0 mRNA, because we detected both mScarlet and the hmNeonGreen mRNA sequences (Figure 2 C).

**Figure 2:**
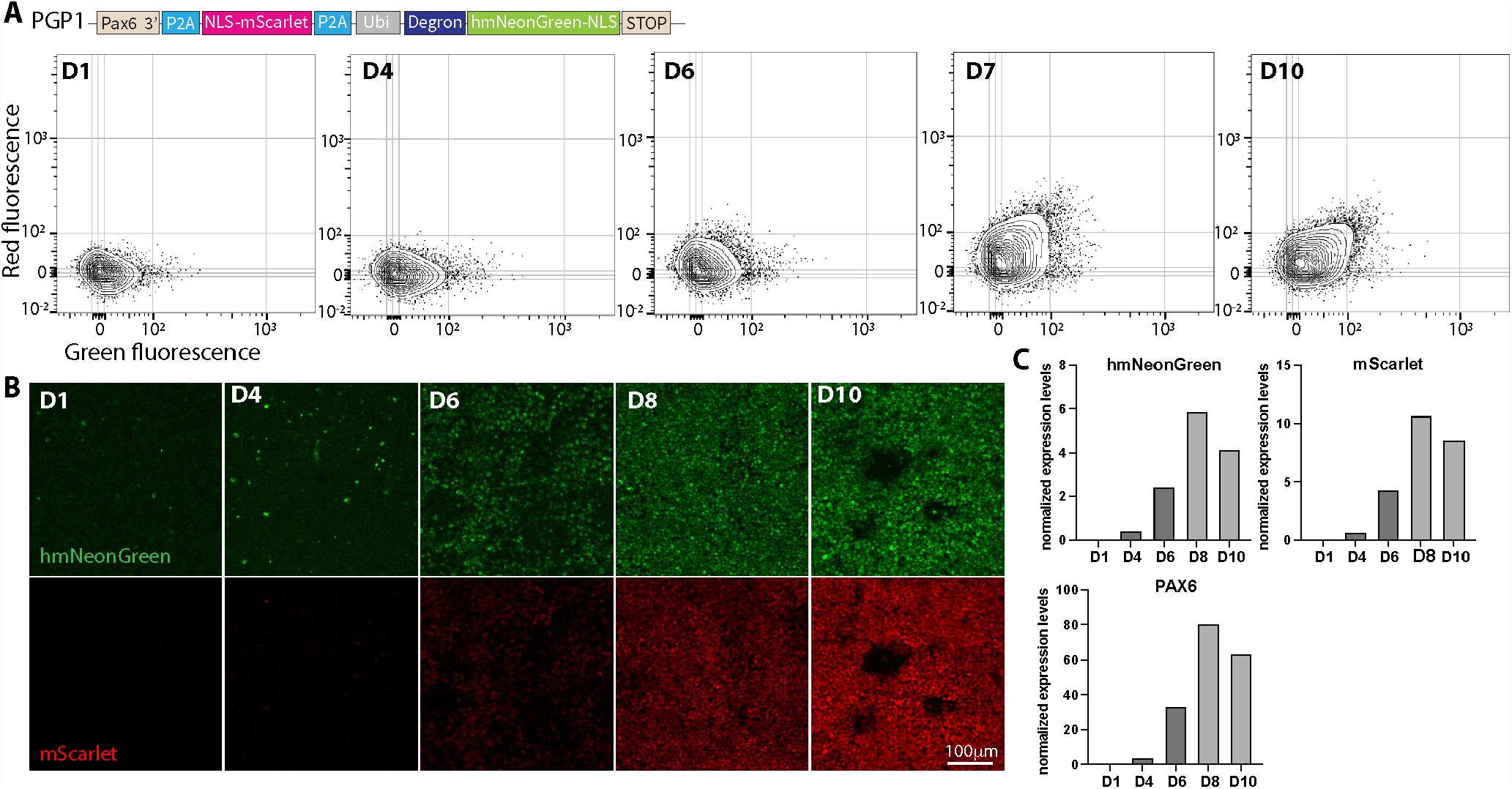
MolTimer1.0 is expressed by human iPSC-derived neural progenitor cells with appropriate multi-day dynamics. A: FACS plots of human iPSC-derived neural progenitor cells expressing MolTimer1.0 from the endogenous PAX6 promoter. PGP1-PAX6-MolTimer1.0 iPSCs were induced at day (D) 0, and neural progenitors were harvested and analyzed daily. Green and red double-positive cells start to appear from D6 onward, with increasing fluorescence intensity until D10. B: Z-projections of confocal imaging stacks of endogenous MolTimer1.0 expression in PGP1-PAX6MolTimer1.0 neural progenitor cells from D1 to D10. Green and red fluorescence is detected from D6 onward, increasing in intensity until D10. C: qPCR analysis of hmNeonGreen, mScarlet, and PAX6 reveals increased mRNA-level expression of both fluorophores together with increased PAX6 expression.

To maintain experimental conditions as directly comparable as possible, and to avoid inherent minor day-today variability of the flow cytometer instrument, flow cytometric analysis and confocal imaging was initially performed on PFA-fixed cells. However, PFA fixation can impact fluorescence intensity of certain FPs, and might make a low hmNeonGreen signal undetectable. Therefore, we repeated the longitudinal fluorescence analysis in live cells, and limited analysis to the times just before we detected fluorescence signal (day 3) to the times when cells were double-positive (day 6). We started the differentiation of all iPSCs on the same day, to avoid variability in the starting cell population, and we analyzed neural progenitor cells on consecutive days, using the same flow cytometer with identical settings. Similar to results in fixed progenitor cells, we first detected a small population of red, mScarlet-expressing cells at day 4 that increased in fluorescence intensity and number. Again, we detected a slight increase in hmNeonGreen fluorescence in the most highly mScarlet-expressing cells. However, we did not detect single fluorescence, hmNeonGreen-only cells (Supplemental Figure 4 A). These results confirm that published maturation times for mScarlet and mNeonGreen, in heterologous expression systems, do not fully reflect maturation times in human neural progenitor cells *in vitro*.

The differentiation protocol we used includes constant dual SMAD inhibition (SMADi) to produce a homogeneous, PAX6-positive, neural progenitor population. Since we did not detect hmNeonGreen-only cells in the PAX6MolTimer1.0 progenitor population, we tested whether a more heterogeneous progenitor pool might enhance identification of hmNeonGreen-only cells.

To achieve a more heterogeneous progenitor pool, we pulse treated PAX6-MolTimer1.0 iPSCs with dual SMAD inhibitors for 24h (D0-D1) and removed the small molecules thereafter. We hypothesized that this short pulse would either lead to no neural induction or to a heterogeneous population of which some cells become neural progenitors and some remain in their undifferentiated pluripotent state. We introduced an additional level of heterogeneity by mixing neural progenitors with matched iPSCs prior to flow cytometric analysis. In these experiments, PAX6MolTimer1.0 progenitors were either pulse treated or continuously treated with dual SMAD inhibitors, then mixed with PAX6-MolTimer1.0 iPSCs (1/20 ratio, progenitors/iPSCs) prior to analysis. Surprisingly, we found that a single, 24h pulse of dual SMAD inhibition led to MolTimer1.0 expression levels comparable to continuous dual SMAD inhibition, and not to the expected higher degree of heterogeneity (Supplemental Figure 4 B). However, our mixing approach succeeded; mixing of PAX6-MolTimer1.0 progenitors with PAX6-MolTimer iPSCs prior to analysis resulted in a heterogeneous population, with only a small distinct fluorescent population in a large non-fluorescent background population. This was particularly apparent on days 5-6, when only a small population displayed hmNeonGreen and mScarlet fluorescence, with most cells not exhibiting any detectable fluorescence (Supplemental Figure 4 C). While these mixing experiments made it easier to identify MolTimer1.0 expressing progenitors, we still did not identify hmNeonGreen-only cells with these experiments.

The MolTimer1.0 design was based on Chrono^27^, but we employed mScarlet instead of dTomato, which has a different maturation time, fluorescence intensity, and fluorescence spectrum, thus might have potentially masked hmNeonGreen fluorescence. Therefore, we generated a PAX6-Chrono iPSC reporter line to test whether Chrono, expressed from the endogenous PAX6 promoter might be utilized to isolate time resolved human neural progenitor cells. We differentiated these PAX6-Chrono iPSCs into neural progenitors using pulsed dual SMAD inhibition, and analyzed them from day 3 to 7 either as a progenitor population or mixed with PAX6-Chrono iPSC prior to analysis. Similar to our observations with MolTimer1.0 expressing progenitors, a small population of mNeonGreen and dTomato double positive cells was first detected from day 4, increasing in number and fluorescence intensity over time. However, we did not detect mNeonGreen-only cells (Supplemental Figure 5). These results demonstrate that MolTimer1.0 and Chrono expressed from the PAX6 promoter do not function as molecular timers in human neural progenitor cells, since we could not capture cells that were single-positive for the fast-maturing mNeonGreen.

MolTimer1.0 and Chrono both contain a destabilized mNeonGreen to theoretically increase the time period over which the FT can report promoter onset. However, we hypothesized that this might also lead to faster degradation of mNeonGreen, making it undetectable. Additionally, mNeonGreen might theoretically mature more slowly in human neural progenitors, leading to its incompatibility with mScarlet or dTomato.

Therefore, we simplified the FT design by removing the degron, and changed hmNeonGreen to sfGFP (MolTimer2.0). We chose mScarlet and sfGFP as the FP pair because both displayed strong expression in neural progenitors (Figure 1 B, 2 A) and because their maturation times are substantially different. We targeted the endogenous PAX6 promoter to generate PAX6-MolTimer2.0 iPSCs (Supplemental Figure 2 B), differentiated those iPSCs into neural progenitors, and longitudinally assayed their fluorescence in PFA fixed cells using fluorescence cytometry and confocal microscopy.

As with all the prior PAX6 reporter lines, we did not detect fluorescence until day 4. From day 4 on, we detected an increase of both sfGFP and mScarlet fluorescence, with both number of detected cells and their intensities increasing roughly linearly until day 10, when all cells showed high level red and green fluorescence. Again, we did not detect sfGFP-only cells with this construct (Figure 3 A). These results were confirmed by confocal microscopy, with no detectable fluorescence until day 4, then a roughly linear increase of both FPs until day 10 (Figure 3 B). QPCR for sfGFP and mScarlet confirmed that the full MolTimer2.0 polycistronic mRNA was present in progenitors, and that its expression increased together with PAX6 during progenitor differentiation. These results exclude simple problems with expression of the MolTimer2.0 construct (Figure 3 C). To again investigate whether there might be negative effects of PFA fixation on the fluorescence signal, we repeated this analysis with live progenitor cells, and followed their fluorescence from day 3 to day 6. Again, we did not detect any sfGFP-only MolTimer2.0-expressing neural progenitors (Supplemental Figure 6 A).

**Figure 3:**
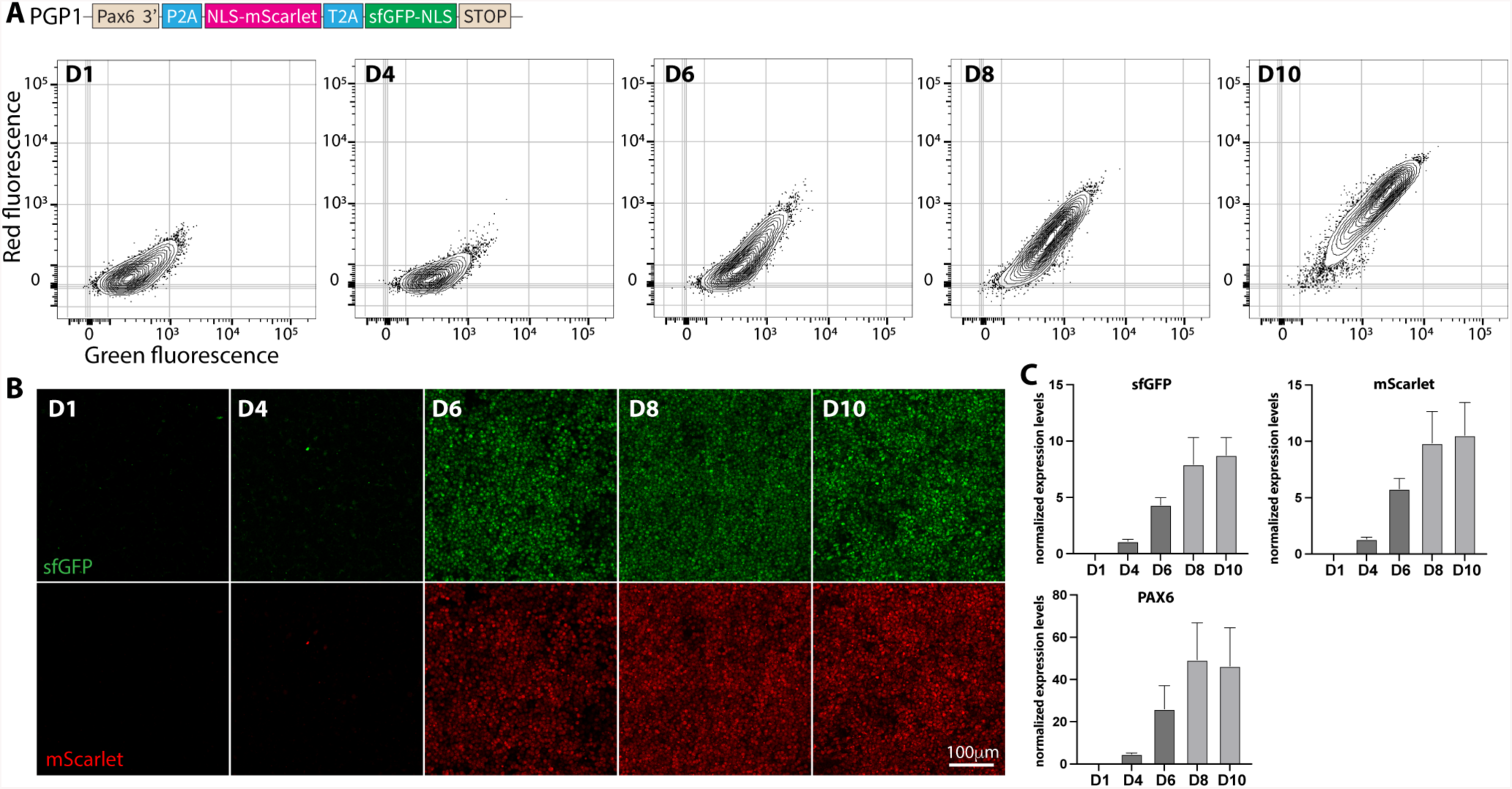
MolTimer2.0 is also expressed by human iPSC-derived neural progenitor cells with appropriate multi-day dynamics. A: FACS plots of human iPSC-derived neural progenitor cells expressing MolTimer2.0 from the endogenous PAX6 promoter. PGP1-PAX6-MolTimer2.0 iPSCs were induced at D0, and neural progenitors were harvested and analyzed daily. Green and red double-positive cells start to appear From D6 onward, with increasing fluorescence intensity until D10. B: Z-projections of confocal imaging stacks of endogenous MolTimer2.0 expression in PGP1-PAX6-MolTimer2.0 neural progenitor cells from D1 to D10. Green and red fluorescence is detected from D6 onward, increasing in intensity until D10. C: qPCR analysis of sfGFP, mScarlet, and PAX6 reveals increasing mRNA-level expression of both fluorophore mRNAs together with increased PAX6 expression.

Previous experiments found that pulsed induction with dual SMAD inhibitors strongly induces MolTimer1.0 expression in neural progenitors (Supplemental Figure 5 B). To again generate a heterogeneous progenitor population, we employed pulsed dual SMAD induction of PAX6MolTimer2.0 iPSC followed by mixing of these induced MolTimer2.0 expressing progenitors with un-induced MolTimer2.0 iPSC. Interestingly, we detected a small sfGFP-only population emerging on day 4 that transitioned into a sfGFP+ and mScarlet+ double-positive population on day 5 and 6 (Supplemental Figure 6 C).

Since an sfGFP-only population was also visible on day 5, we asked whether these cells might represent an early, immature progenitor population that just started to express PAX6 and is distinct from slightly more mature doublepositive progenitors. We tested this hypothesis by FACS isolation and subsequent qPCR analysis for PAX6 and sfGFP expression on day 5. We pulse-differentiated PAX6MolTimer2.0 iPSC into neural progenitors, mixed them with un-induced PAX6-MolTimer2.0 iPSC (1/20 ratio, progenitors/iPSCs) and isolated four distinct populations based on ratiometric green and red fluorescence (Figure 4 A, B). One population consisted of non-fluorescent cells, expected to contain mostly PAX6-negative iPSCs. As a second population, we collected weakly GFP+ cells, expected to consist of progenitors that just started to express MolTimer2.0, with low PAX6 expression levels. The third population consisted of sfGFP/mScarlet double-positive cells, expected to contain more mature progenitors with moderate PAX6 levels. The fourth population consisted of high level expressing sfGFP/mScarlet double-positive cells, expected to contain the most mature, highly PAX6-expressing progenitor population (Figure 4 C). Of note, we detected roughly equally high levels of PAX6 and sfGFP expression in the sfGFP/mScarlet double-positive populations. In contrast to our expectations, we did not detect PAX6 or sfGFP expression in the sfGFP-only population, indicating that this population might mainly consist of iPSC, and not a distinct, early PAX6-expressing progenitor population (Figure 4 D).

**Figure 4:**
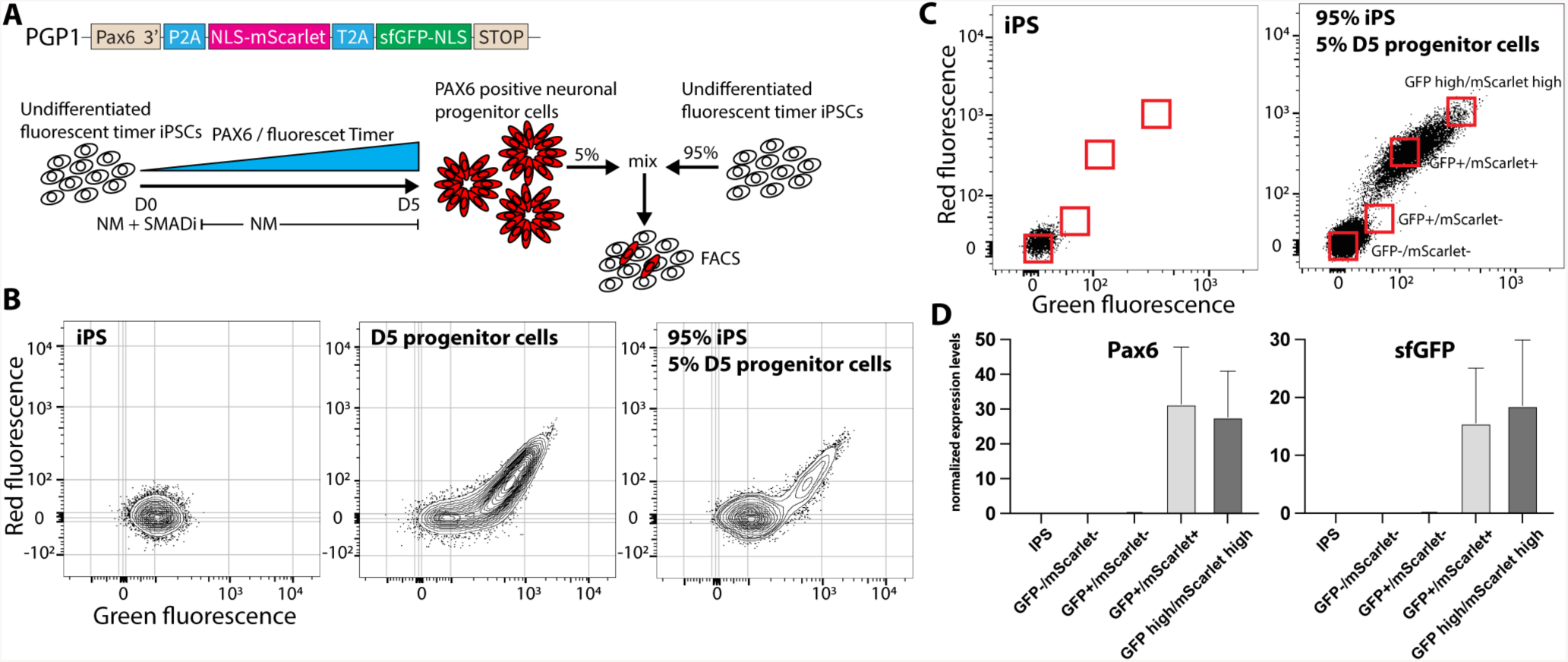
FACS isolation and qPCR of MolTimer2.0 expressing neural progenitor cells. A: Schematic of the mixing and FACS isolation of MolTimer2.0 expressing neural progenitor cells. PGP1-PAX6-MolTimer2.0 neural progenitors were mixed with undifferentiated PGP1-PAX6MolTimer2.0 iPSC prior to FACS analysis. B: FACS plots of PGP1-PAX6-MolTimer2.0 iPSC (left), D5 PGP1-PAX6-MolTimer2.0 neural progenitor cells (middle), and the 1:20 mix of D5 PGP1-PAX6-MolTimer2.0 progenitor cells and undifferentiated PGP1-PAX6-MolTimer2.0 iPSCs used for isolation and qPCR analysis (right). C: Gating strategy to isolate four distinct populations of MolTimer2.0 expressing neural progenitor cells: nonfluorescent GFP-/mScarlet-, green-only GFP+/mScarlet-, dual fluorescent GFP+/mScarlet+, and high level dual fluorecent GFP+/mScarlet+. D: qPCR analysis of PAX6 and sfGFP expression. PAX6 and sfGFP are only detected in the low level GFP+/mScarlet+ and high level GFP+/mScarlet+ populations.

## Discussion

We investigated the feasibility of multiple, progressively refined fluorescent timer (FT) proteins to report “developmental time” after onset of expression of a developmentally indicative transcription factor (TF), as opposed to chronologic time. In particular, we tested two new designs, MolTimer1.0 and MolTimer2.0, and the Chrono construct published in 2019 by Gehart et al.^27^, during neural differentiation of human iPSCs into neural progenitor cells. Ideally, multiple, spectrally distinct FT proteins would be expressed from the promoter of such a fate-determining TF. The sequential expression of those FTs, and their ratiometric use in FACS, would operate in a sense like a chronometer, enabling isolation of distinct, time resolved cellular populations during differentiation of human pluripotent stem cells into subtype-specific neurons or other cell types.

As a first step toward this goal of a sequential FT protein expression system, we engineered five human iPSC lines to express molecularly distinct FTs from the endogenous PAX6 promoter. These FTs ranged from a single FT that changes color over time, SlowFT, to constructs of various levels of complexity with distinct green and red fluorescent proteins, including the recently published Chrono^27^. We found that four out of these five FT constructs lead to strong and readily detectable fluorescence upon neuronal induction, with their expression levels increasing with PAX6 expression. In these four FT constructs, we combined a fast-maturing green fluorescent protein with a slow-maturing red fluorescent protein. We predicted that the difference in maturation times would enable ratiometric green-to-red fluorescence measurements, thus isolation of distinct neural progenitor cells. In contrast to these predictions and results in pilot HEK 293T cells, we found that fluorescence of both fluorophores increased in tandem in a roughly linear fashion when expressed from the PAX6 promoter. In contrast, 293T cells first exhibited green fluorescence, then slowly became double-positive, faithfully timing promoter activation as we predicted with these rationally designed constructs.

Our results raise the question of why the FT protein constructs faithfully time promoter onset in 293T cells, or with Chrono in mouse enteroendocrine progenitor cells, but not in human iPSC-derived neural progenitor cells. First, expression levels are different between 293T and neural progenitor cells. When the PAX6 promoter becomes active, initial FT protein levels are low, so green fluorescence is likely below detection threshold. However, on average, the PAX6 promoter is one of the strongest promoters in neural progenitor cells^39^. Even with the relatively strong expression of the PAX6 promoter, we detected much higher fluorescence when we heterotopically expressed FT proteins in 293T cells. A second possible explanation is sampling frequency. We assayed cells every 24h, and might have missed a very short phase when only the fast-maturing FP was detectable in maturing neural progenitor cells. However, in 293T cells, we detected the first green cells 6h after promoter activation, and could still detect GFP-only cells 24h later, making this explanation less likely.

Potentially more likely, fluorescent proteins might mature differently in distinct cell types. Except for DsRed2, we always detected both green and red fluorescence in neural progenitor cells, indicating correct processing of both proteins. However, it remains unclear how efficiently FT proteins mature in neural progenitor cells after they are expressed and translated. We carefully selected fluorescent protein pairs based on their maturation times and selected spectrally distinct variants with large differences in their known maturation times. However, those values might be quite different in previously unstudied neural progenitor cells compared to published values derived in pure protein extracts, heterotopic expression systems, or *E*.*coli*^40^.

Further, it is known that distinct promoters have different activation and expression kinetics^41^. Human neurogenesis *in vivo* and *in vitro* is a lengthy process that occurs over weeks and months compared to days in mice. The PAX6 promoter is one of the first promoters activated when human iPSC exit their pluripotent state and become committed toward neural fate. During this process, PAX6 slowly increases in neural progenitor cells over a long period of time, which also leads to slow, steady increase of FT expression in dividing neural progenitors. This is quite different from pulsed and transient FT expression in 293T cells or in mouse enteroendocrine cells that differentiate from a common progenitor into mature cells in 24h. In a pulsed system, a limited amount of an FT is made relatively synchronously, then matures, thus operating effectively as a developmental “chronometer” reporting promoter activation. Our experiments indicate that, in striking and limiting contrast, FT expression from the PAX6 promoter in neural progenitor cells slowly increases. Thus, it is likely that each individual cell contains a mixture of an individual FT at a range of protein maturation stages. This might likely explain our experimental results of lack of detectable ratiometric distinction between two FT proteins after early, possibly weak fluorescence following PAX6 promoter activation.

Finally, there are differences between expression systems. Early phases of *in vitro* neural differentiation from iPSCs appear largely synchronized, with most cells expressing PAX6 at similar levels. This would likely make it difficult to identify subtly developmentally distinct cells based on small fluorescence differences between cells. This is in sharp contrast to the mouse enteroendocrine system, in which only a small population starts to express a promoter, thus FT protein, at distinct times, thus enabling isolation of time-resolved cells from a largely dark, non-FT-expressing background. Though we predicted this difference between mouse and human systems, and designed our FT constructs and experiments to overcome this by mixing FT-expressing neural progenitors with matched iPSCs prior to analysis, this strategy was not sufficient. While these design steps made it possible to identify FT-expressing cells, we were unsuccessful in differentially isolating cells that had just started to express PAX6 vs. cells that were developmentally more mature. These results reinforce the presence of quite homogeneous and developmentally synchronized neural progenitors. Using a different promoter, only active in a select population of cells, and ideally only transiently, might overcome this limitation.

Taken together, our sequential design and testing of a range of five knock-in hPSC lines that express five distinct single and dual fluorescent timer proteins from the endogenous paired box 6 (PAX6) promoter demonstrates that none of the tested FP/FT constructs were successful in identification and isolation of distinct progenitor populations *in vitro* based on ratiometric fluorescence and FACS. Such isolation of distinct, developmentally synchronized cortical progenitors *in vitro* would enable direct investigation of developmental trajectories of transcriptional regulation. While these dual FP or FT systems faithfully reported chronologic time when expressed from a strong inducible promoter in 293T cells, none of the tested FP/FT constructs followed the same fluorescence kinetics in developing human neural progenitor cells. Our comparison to 293T cells, and the use of a published FT, demonstrates that FT proteins need to be carefully selected and tested for specific cell types and differentiation systems. This work highlights unique and often surprising expression kinetics and regulation in specific cell types differentiating from hPSCs.

## Supporting information

qPCR primer list

## Author contributions

M.P., S.S., and J.D.M conceived the overall project and experiments;

M.P. and S.S. designed initial experiments, M.P. and J.D.M. designed later experiments, and M.P. performed the experiments; M.P. and J.D.M. analyzed and interpreted the data, integrated the findings, and wrote and edited the manuscript. All authors contributed to discussions and manuscript editing.

## Acknowledgements

We thank Holly McKee and Karen Wang for their excellent technical support; members of the Macklis laboratory for scientific discussions and helpful suggestions; Joyce LaVecchio and Nema Kheradmand of the HSCRB-HSCI Flow Cytometry Core; the Harvard Center for Biological Imaging for infrastructure and support. This work was supported by the following grants to J.D.M.: Paul G. Allen Frontiers Group Allen Distinguished Investigator award #11855; National Institutes of Health OD DP1 NS106665; additional infrastructure support from NIH grants NS104055 NS045523, and NS049553; and the Max and Anne Wien Professor of Life Sciences fund.

## Materials and Methods

### Cell culture

All cell lines were maintained at 37°C at 5% CO2 in a humidified incubator.

The human induced pluripotent stem cell line PGP1 (Personal Genome Project 1) was obtained from the laboratory of G.Church42. iPSCs were grown in mTeSR+ medium (StemCell Technologies, 100-0276) supplemented with penicillin/streptomycin (Pen/Strep, 1% vol/vol, ThermoFisher, 15140122) on Geltrex-coated coated cell culture dishes (ThermoFisher A1413302). 293T cells were obtained from ATCC and grown in DMEM (ThermoFisher, 10566016) supplemented with FBS (10% vol/vol, ThermoFisher, 10438026) and Pen/Strep (1% vol/vol).

### Generation of doxycycline (DOX) inducible, knock-in fluorescent timer (FT) 293T cells

DOX-inducible 293T-FT cells were generated using the piggyBack transposon system. The FT construct was cloned into a piggyBack transposon targeting vector containing a tetracycline-inducible expression cassette and a puromycin selection cassette flanked by piggyBac transposase specific inverted terminal repeats. 293T cells were grown to 60% confluency in one well of a 6 well plate, and co-transfected with 1ug transposon targeting plasmid and 1ug transposase plasmid using Lipofectamin3000 (ThermoFisher, L3000015) following the manufacturer’s protocol. Medium was changed 24h later, and cells were incubated for additional 48h. After 48h, puromycin (1ug/ml, ThermoFisher, A1113803) was added to the medium. 48h later, cells were harvested with TrypsinEDTA (ThermoFisher, 25200056) to generate a single cell solution, and single clones were collected into 96 well plates using FACS for genotyping. Positive clones were frozen in complete DMEM medium supplemented with DMSO (20% vol/vol, SantaCruz, sc-358801).

### DOX induction of FT in 293T cells

DOX inducible FT 293T cells were grown in complete medium to reach 50% confluency. The FT was induced by adding doxycycline (500ng/ml, Sigma, D9891) to the complete medium. 24h later, medium was changed to fresh medium without doxycycline. Cells were harvested with trypsin-EDTA and prepared for FACS analysis.

### Generation of PAX6-fluorescent timer lines

We used CRISPR/Cas9 editing to integrate one of a series of iterative FT constructs downstream of the endogenous PAX6 promoter. The donor plasmid contained (from 5’ to 3’) the 3’end of the PAX6 gene without the stop codon, a P2A peptide, and one of this set of iterative FT constructs (“SlowFT”, “TandemFP”, “MolTimer 1.0”, “MolTimer 2.0”, “Chrono”), and the DNA sequence directly downstream of the PAX6 stop codon. TandemFP consists of a fusion of DsRed2 and sfGFP; MolTimer 1.0 consists of NLSmScarlet-P2A-Ubiquitin-Degron-humanNeonGreen-NLS and MolTimer 2.0 consists of NLS-mScarlet-T2A-sfGFP-NLS. The Chrono plasmid27 was generously provided by Professor Hans Clevers, University Utrecht). The Cas9-guideRNA plasmid pSpCas9(BB)-2A-GFP (Addgene, 48138) was obtained from Addgene, and the PAX6 guideRNA (5’ GGCCAGTATTGAGACATATC 3’) was cloned downstream of the human U6 promoter.

To integrate one of a set of FT constructs downstream of the PAX6 promoter, iPSCs were first disassociated with Accutase (Innovative Cell Technologies, AT104) into a single cell suspension. Nucleofection was performed with an Amaxa 4D-Nucleofector System (Lonza) using the P3 Primary Cell nucleofector kit (Lonza, V4XP-3012). 8×105 cells were resuspended in 100µl P3 solution, combined with 3mg donor plasmid and 2mg Cas9-guideRNA plasmid, and nucleofected using the CA-137 program. After nucleofection, cells were plated on Geltrex-coated 10cm plates in mTeSR+ supplemented with CloneR (StemCell Technologies, 05888). A full medium change was performed 48h post nucleofection with fresh mTesR+ supplemented with CloneR. Four days after nucleofection, the medium was changed to fresh mTesR+ containing 0.3ug/ml puromycin. Medium changes were performed daily from day 4 onwards. On day 6 the medium was changed to fresh mTeSR+ without puromycin, and cells were grown until clones were manually picked into 96 well plates for genotyping. Genomic DNA was isolated with the QuickExtract DNA Extraction Solution (Lucigen, QE09050), and correct PAX6 integration of the appropriate FT construct was confirmed by PCR. The full FT cassette was Sanger sequenced to confirm correct genomic integration into the 3’ end of the PAX6 gene, and clones were banked in mFreSR freezing medium (StemCell Technologies, 05854).

### Cortical differentiation

Neuronal induction was performed as described previously43. In brief, iPSCs were disassociated with Accutase to generate a single cell suspension, and 2.8×105 cells/cm2 were plated on Geltrex-coated plates in mTesR+ supplemented with ROCK inhibitor Y-27632 (10uM, StemCell Technologies, 72302). Neuronal induction was started 24h later (day 0) by changing mTesR+ medium to neuronal induction medium consisting of neuronal maintenance medium (NMM) supplemented with SB43152 (10µM, Selleckchem, S1067) and Dorsomorphin (DM, 10µM, Stemcell Technologies, 72102). The medium was changed daily until day 12. For pulsing experiments DM and SB43152 was omitted from the NMM from day one onwards.

NMM consisted of DMEM:F12 (50% vol/vol, ThermoFisher, 10565018), Neurobasal medium (50% vol/vol ThermoFisher, 21103049), insulin (0.03% vol/vol, Sigma, I9278), 2-mercaptoethanol (0.1% vol/vol, ThermoFisher, 31350010), non-essential amino acids (0.5% vol/vol, ThermoFischer, 11140050), Sodium Pyruvate (0.5% vol/vol Sigma, S8636), Pen/Strep (0.5% vol/vol, ThermoFisher, 15140122), N-2 supplement (0.5% vol/vol, ThermoFisher, 17502048), B-27 supplement (1% vol/vol ThermoFisher, 17504044) and Glutamax (0.5% vol/vol ThermoFisher, 35050061).

### qPCR analysis

Cells were harvested with Accutase and washed twice with PBS. RNA was isolated using the RNeasy Plus Micro kit (Quiagen, 74034), and cDNA was synthesized with the SuperScript IV First-Strand Synthesis system (ThermoFisher, 18091050) following the manufacturer’s protocols. PCR targets were amplified with the PowerUp SYBR Green Master Mix (ThermoFisher, A25741) using the primer sequences in the “qPCRPrimer” supplementary table.

### Western blot analysis

Cells were harvested with Accutase and washed twice with PBS. Whole cell protein extraction was performed by lysing the cell pellet with Cell Lysis buffer (ThermoFisher, FNN0011) supplemented with protease inhibitor cocktail (ThermoFisher, 78442) following the manufacturer’s protocol. Western blot analysis was performed with the following antibodies: β-actin (1:10,000 Sigma, A5441), GFP (1:1,000, ThermoFisher, A11122), DsRed2 (1:200 SantaCruz, sc-101526), PAX6 (1:500 ThermoFisher, AD2.38), RPL22 (1:1,000 SantaCruz, sc-373993), mNeonGreen (1:1,000 CellSignallingTechnologies, 53061), mScarlet (1:1,000, Rockland 600-401-379)

### Confocal imaging

Cells were differentiated on 8 well µ-Slides (Ibidi, 80826). Cells were washed twice with PBS, and fixed with PFA (4% w/vol, ThermoFisher, 28906) for 10min at room temperature (RT). PFA was removed, and cells were washed twice with PBS and stored in PBS at 4°C. On the imaging day, the PBS was removed, and cells were permeabilized by incubation with PBS containing Triton-X100 (0.3% vol/vol, Sigma Aldrich X-100) for 5min at RT. Nuclei were stained with 4’,6-Diamidino-2-Phenylindole (DAPI, 5µg/ml, DAPI, ThermoFisher D1306) in PBS for 5 min at RT, followed by a PBS wash. Cells were imaged on an inverted Zeiss LSM 880 confocal microscope. All images were processed with Fiji44.

### Live cell imaging

Cells were differentiated on 8 well µ-Slides and imaged for 24h at a 20min time interval on a Zeiss Cell Discoverer7 live-cell-imaging system.

### Flow cytometry analysis

For fixed FACS analysis, cells were harvested with Accutase to generate a single cell suspension and washed twice with PBS. Cells were incubated for 5min with PFA (4% w/vol), washed twice with PBS, and stored at 4°C. Cells were filtered through a 35µm cell strainer (Corning, 352235) prior to analysis. For live cell FACS analysis, cells were harvested with Accutase, washed twice with PBS, and filtered through a 35µm cell strainer. Cells were kept on ice and incubated with SYTOX Blue Dead Cell Stain (ThermoFisher, S34857) prior to FACS analysis. All samples were analyzed on a LSRII (BD Bioscience) or FACSymphony A5 (BD Bioscience), cytometer and plots were prepared using FlowJo (BD Bioscience).

## Supplemental Figures

**Supplemental Figure 1:**
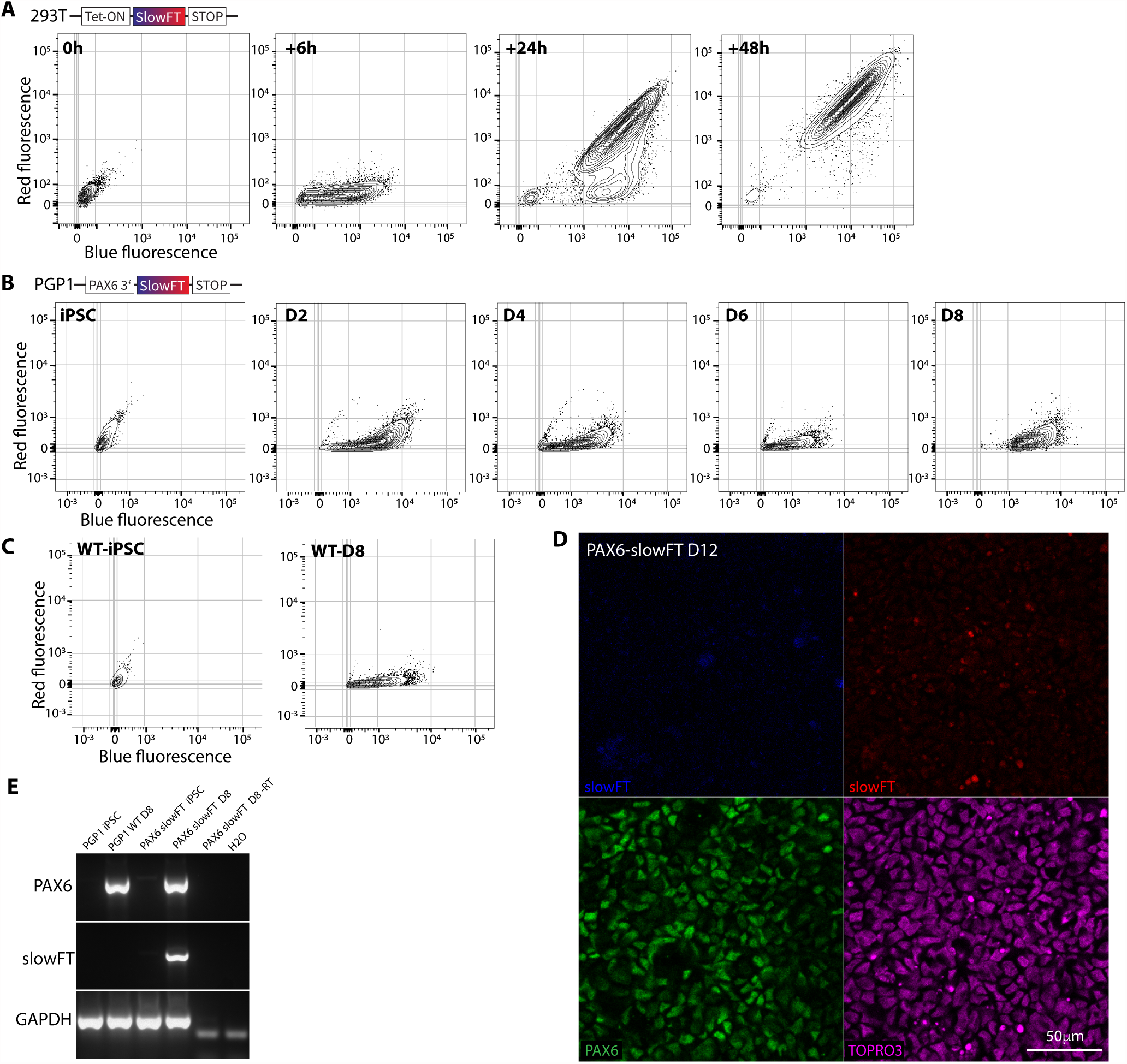
SlowFT expression has distinct dynamics in 293T cells vs. neural progenitor cells. A: FACS plots of 293T cell expressing SlowFT from the Tet-ON promoter. Expression was induced by adding Dox to the cell culture medium. Cells were harvested and analyzed 6h, 24h, and 48h later. Blue fluorescence strongly increases by 6h after induction, followed by most cells displaying both blue and red fluorescence at 24h and 48h. B: FACS plots of PGP-1 iPSC-derived neural progenitor cells expressing SlowFT from the endogenous PAX6 promoter. PGP1PAX6-SlowFT iPSCs were induced to undergo neural differentiation at D0. Neural progenitors were harvested and analyzed daily for eight days. Blue fluorescence strongly increased by D2, without similar increase of red fluorescence through D8. C: FACS plots of wild type (WT) PGP1 iPSCs and WT D8 neural progenitor cells. Note the strong increase of blue fluorescence in WT cells that do not express SlowFT. D: Confocal imaging stack of PAX6-slowFT neural progenitor cells at day 12. No blue or red cellular fluorescence is detected. Bright red signal is fluorescent debris outside of cell bodies. E: RT-PCR of WT and PAX6-SlowFT iPSCs and neural progenitor cells. PAX6 is highly expressed by both WT and PAX6-SlowFT neural progenitor cells. PAX6-SlowFT mRNA is only detected in PAX6-SlowFT neural progenitor cells.

**Supplemental Figure 2:**
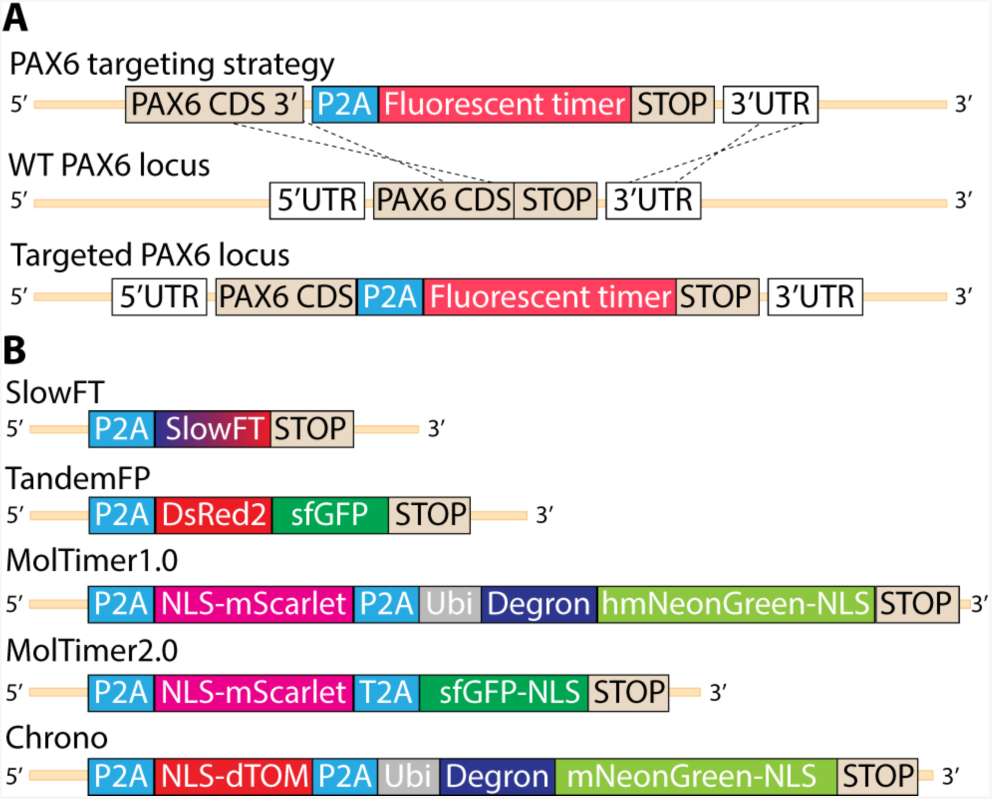
Targeting strategy enables expression of fluorescent timer constructs under control of the human PAX6 promotor. A: Targeting strategy for the integration of fluorescent timer constructs and their expression from the endogenous PAX6 promoter. The 3’ stop codon of the PAX6 gene was removed, and each fluorescent timer construct together with a P2A site was integrated in frame with the PAX6 gene. B: Fluorescent timer constructs used in this study. SlowFT consists of a single fluorescent timer molecule that matures and changes color from blue to red. TandemFP is a fusion of the red FP DsRed2 and the green FP sfGFP. MolTimer1.0 consists of a nuclear localized (NLS) mScarlet and NLS humanized hmNeonGreen separated by a P2A site. hmNeonGreen is destabilized with a Degron. MolTimer2.0 consists of NLS mScarlet and NLS sfGFP separated by a T2A site. Chrono27 consists of NLS dTom and a destabilized NLS mNeonGreen separated with a P2A site; it was used as a comparison strategy published following development and application in mice.

**Supplemental Figure 3:**
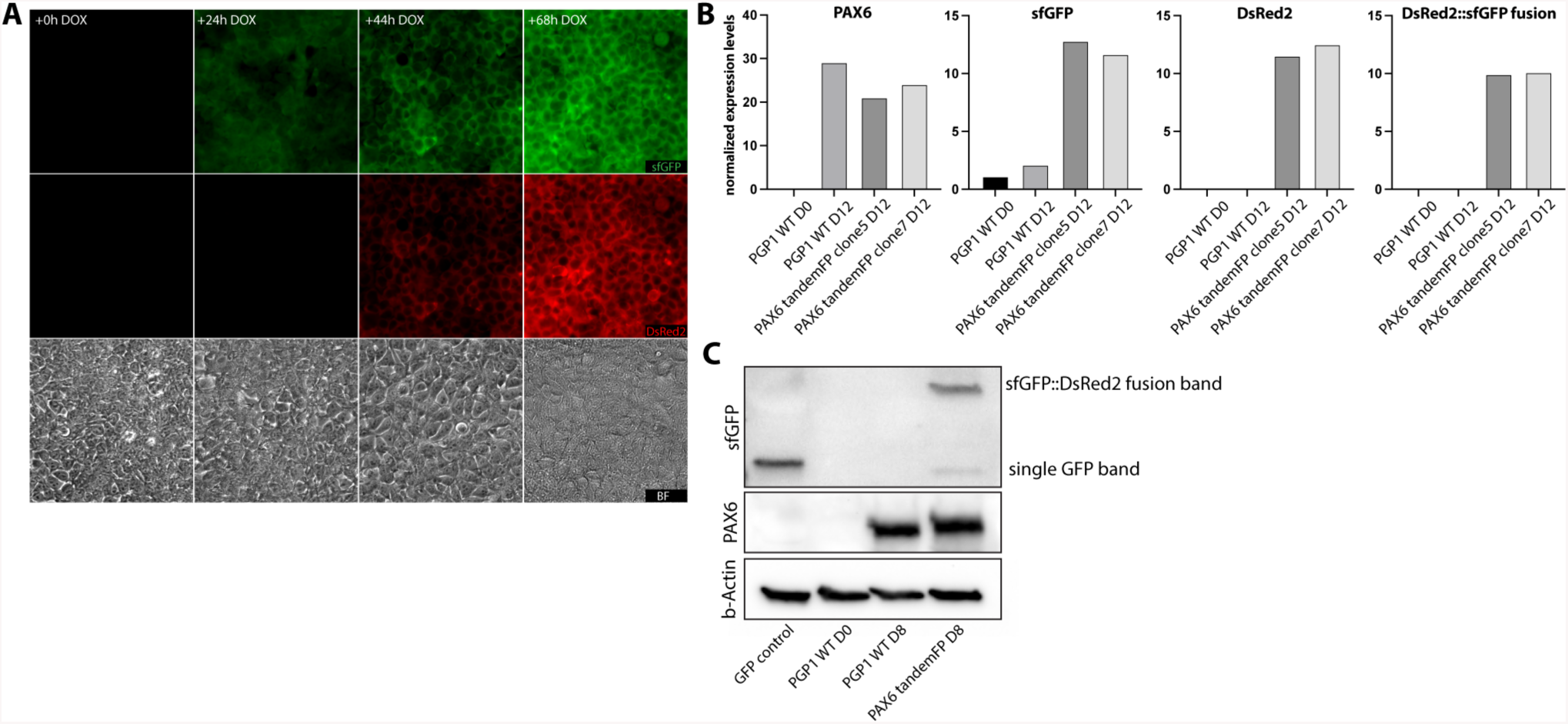
TandemFP is expressed appropriately by both 293T cells and human iPSC-derived neural progenitor cells at both RNA and protein levels. A: Fluorescence and brightfield images of TandemFP-expressing 293T cells after Dox induction. Green fluorescence is detected by 24h after induction. Green and red double-positive cells appear by 24h and increase in number and fluorescence intensity thereafter. B: qPCR of WT PGP1 hiPSCs and PGP1-PAX6-tandemFP expressing hiPSC and neural progenitor cells. The sfGFP, DsRed, and sfGFP::DsRed2 fusion mRNAs increase along with PAX6 in PGP1-PAX6-tandemFP neural progenitor cells. C: WB for PAX6 and sfGFP of WT PGP1 and PGP1PAX6-tandemFP progenitor cells. The sfGFP::DsRed2 fusion band is detected in PGP1-PAX6-tandemFP neural progenitor cells, indicating both expression and translation of the TandemFP fusion protein.

**Supplemental Figure 4:**
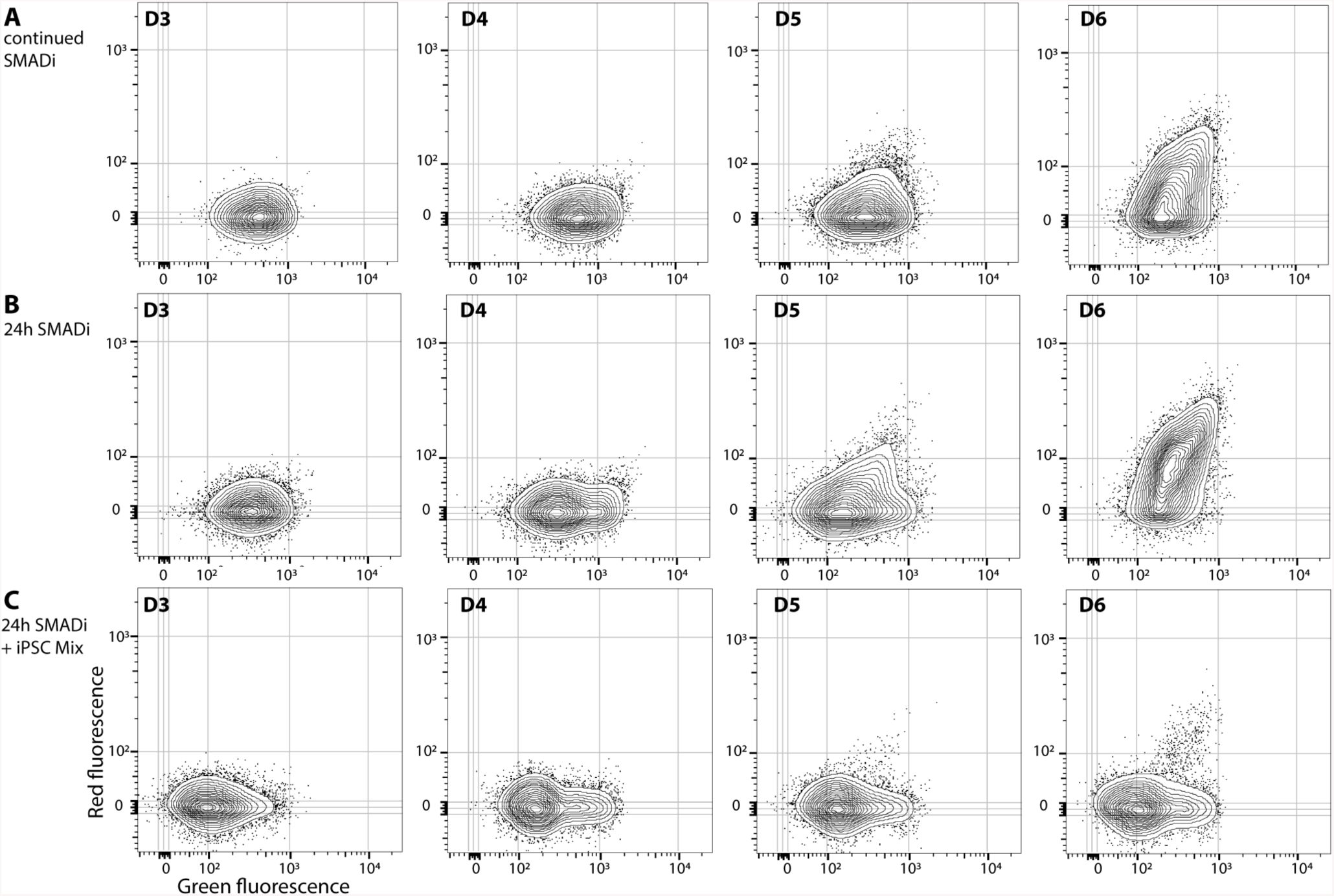
Dynamic live cell FACS analysis of MolTimer1.0 expression by human iPSC-derived neural progenitor cells indicates strong red fluorescence but only weak green fluorescence, and detection of fluorescent PAX6-expressing cells against an overall background of non-fluorescent iPSCs. A: FACS plots of live PGP1-PAX6-MolTimer1.0 neural progenitor cells. Neural induction was induced with continuous dual SMAD inhibition (SMADi). Red fluorescent cells appear by D4, and increase in number and intensity through D6. Weakly green fluorescent cells appear by D5. B: FACS plots of PGP1-PAX6-MolTimer1.0 neural progenitor cells that were induced by a pulse of SMADi for 24h from D0 to D1, yielding very similar results as continuous SMADi induction. C: FACS plots of MolTimer1.0expressing neural progenitors that were mixed at a 1:20 ratio with uninduced PGP1-PAX6-MolTimer1.0 iPSC prior to analysis. Red fluorescent cells can be detected by D5 against a background of >95% non-fluorescent iPSCs.

**Supplemental Figure 5:**
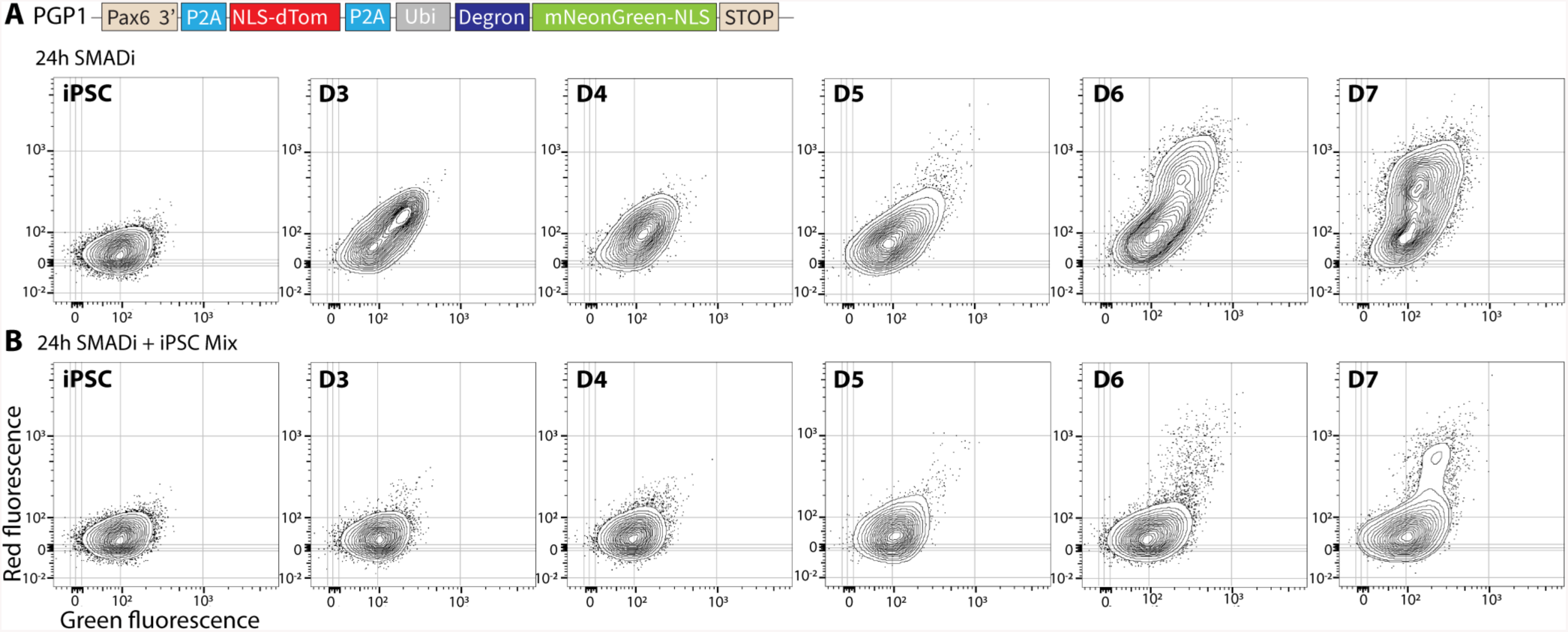
Dynamic live cell FACS analysis of Chrono27 expression by human iPSC-derived neural progenitor cells indicates dual fluorescence expression, and detection of PAX6-expressing cells against a background of overall non-fluorescent iPSCs. A: FACS plots of live PGP1-PAX6-Chrono neural progenitor cells. Neural differentiation was induced for 24h with SMADi. Red and green double-positive cells can be detected by D3, with increased numbers of cells and fluorescence intensity through D6-7. B: FACS plots of undifferentiated PGP1-PAX6-Chrono iPSCs and Chrono-expressing neural progenitors mixed at a 1:20 ratio with undifferentiated PGP1-PAX6-Chrono iPSC prior to analysis, indicating that red and green double-positive cells can be detected by D3 against a background of >95% non-fluorescent hiPSCs. No green or red signal is detected in undifferentiated PGP1-PAX6-Chrono iPSCs.

**Supplemental Figure 6:**
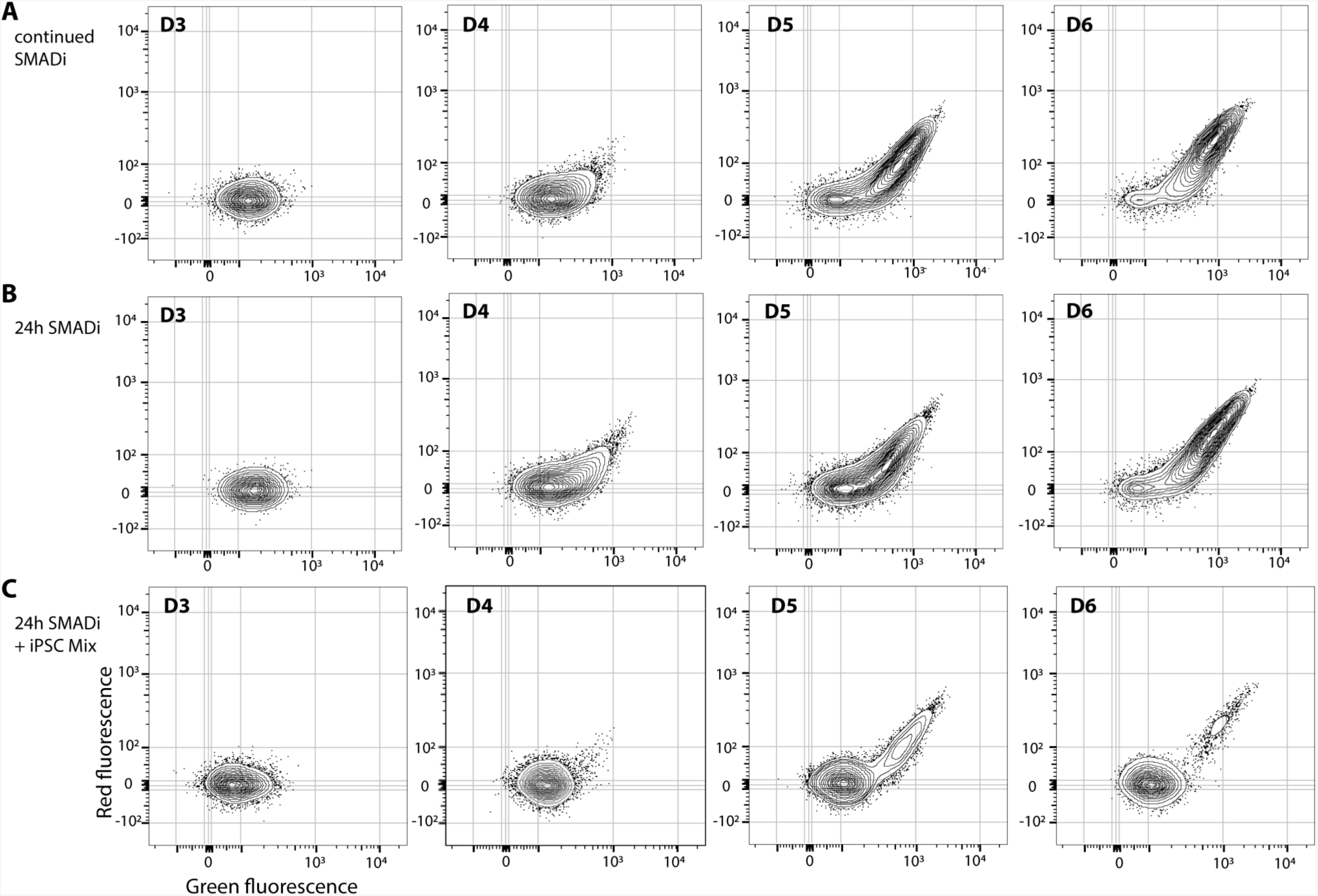
Dynamic live cell FACS analysis of MolTimer2.0 expression by human iPSC-derived neural progenitor cells indicates dual fluorescence expression, and detection of PAX6-expressing neural progenitors against a background of >95% non-fluorescent iPSCs. A: FACS plots of live PGP1-PAX6-MolTimer2.0 hiPSC-derived neural progenitor cells. Neural differentiation was induced with continuous SMADi. Red and green double-positive cells can be detected by D4, with increasing number and fluorescence intensity through D6. B: FACS plots of MolTimer2.0 neural progenitor cells that were induced by a 24h pulse of SMADi from D0 to D1. C: FACS plots of MolTimer2.0-expressing neural progenitors that were mixed at a 1:20 ratio with undifferentiated PGP1-PAX6-MolTimer2.0 hiPSCs prior to analysis. Red and green double-positive fluorescent cells can be detected by D4 against a background of >95% non-fluorescent hiPSCs.

